# USP14 regulates pS129 α-synuclein levels and oxidative stress in human SH-SY5Y dopaminergic cells

**DOI:** 10.1101/2024.05.09.592905

**Authors:** Vignesh Srinivasan, Rabah Soliymani, Larisa Ivanova, Ove Eriksson, Nina Peitsaro, Maciej Lalowski, Mati Karelson, Dan Lindholm

**Author notes:** Corresponding author: Dan Lindholm, Medicum, Department of Biochemistry and Developmental Biology, University of Helsinki, Finland.

## Abstract

Ubiquitin specific protease-14 (USP14) is critical for controlling protein homeostasis disturbed in human disorders like Parkinsońs disease (PD). Here we investigated the role of USP14 in regulating proteasome and autophagy pathways, and their influence on α-synuclein (α-syn) degradation. Data showed that α-syn and phosphorylated serine129 α-syn (pS129 α-syn) were elevated in *USP14* gene-deleted SH-SY5Y dopaminergic cells with concomitant reduction in proteasome activity. Inhibition of proteasomes using MG132 particularly elevated pS129 α-syn in these cells, but the levels were not influenced by inhibiting autophagy using chloroquine. In contrast, autophagy and the CLEAR (Coordinated Lysosomal Expression and Regulation) pathways were elevated in USP14 lacking cells with an upregulation of the transcription factor TFEB. USP14-ablated cells also exhibited increases in reactive oxidative species (ROS) and elongation of mitochondria. The addition of N-Acetylcysteine amide (NACA) to counteract oxidative stress, reduced pS129 α-syn and α-syn levels in USP14 deficient cells. Phospho-proteomic analyses revealed that USP14 is phosphorylated at S143 affecting its function and structure as shown by molecular modeling, and protein interaction studies. Re-expression of wild-type and the phospho-mimetic S143D-USP14 mutant decreased ROS, pS129 α-syn, and α-syn in USP14 lacking cells. These results demonstrate that pS129 α-syn levels are sensitive to oxidative stress in SH-SY5Y dopaminergic cells. USP14 by stimulating the proteasome activity and reducing oxidative stress is a promising factor for targeting α-syn and its pathogenic variants in PD.

## Introduction

Disturbed protein quality control is associated with several human diseases, including cancer and neurodegenerative disorders. Protein-folding by molecular chaperones, degradation via the ubiquitin-proteasome system (UPS), and autophagy-lysosome pathway (ALP) contribute to optimal proteostasis ^1, 2, 3^. UPS and ALP can act in conjunction to control cellular protein levels and intra-cellular organelle homeostasis in health and disease ^4, 5, 6^. However, the precise mechanisms by which these systems are interconnected in the context of cellular physiology, are largely unknown.

Parkinson’s disease (PD) is associated with an increase in soluble and aggregation-prone α-syn levels due to defects in protein degradation, and folding mechanisms ^7, 8^. α-Syn accumulates in intracellular aggregates termed Lewy bodies that form within PD-susceptible midbrain dopaminergic neurons. This is accompanied disturbed protein handling, mitochondria defects, increased oxidative stress-associated damage, and ER stress ^9, 10^. The mechanisms by which misfolded α-syn are elevated in PD are not fully understood but defects in proteasome and autophagy are proposed to contribute to the disease.

The deubiquitinating enzyme USP14 is known to influence the 26S proteasomes by different mechanisms ^11, 12, 13, 14, 15, 16, 17, 18, 19, 20^, and is reversibly associated with the 19S regulatory particle (RP) subunits, PSMD2, and PSMC2 ^11, 14, 17, 18^. In yeast, ubiquitinated proteins upon binding to USP14, activate the 26S proteasome by inducing sequential changes to the 19S RP, facilitating the entry of unfolded substrates into the proteasome core particle (CP), after removal of the polyubiquitin chains by USP14 and Rpn11^18, 19, 21^. The binding of USP14 to the proteasomes increases its activity about 800-fold ^11, 15^. Binding of the UBL domain in USP14, or other UBL-domain containing proteins, such as Rad23, activates the proteasome ^18, 22^. USP14 further removes ubiquitin chains on substrates, antagonizes protein degradation and stabilize their levels ^16, 23^.

Physiologically, USP14 influences cellular pathways such as autophagy and ER stress ^24, 25^. USP14 is linked to different diseases, including cancer ^26, 27, 28, 29, 30^, but its role in neurodegenerative disorders has been less studied. We show here that deletion of *USP14* in SH-SY5Y human dopaminergic cells elevates α-syn, and its S129 phosphorylated form (pS129 α-syn) associated with PD ^31, 32, 33, 34, 35^ that was linked to reduced proteasome activity and elevated reactive oxygen species (ROS), Notably, TFEB and autophagy were upregulated in the USP14-deleted cells with little effect on α-syn clearance. Phosphoproteomic analyses revealed that USP14 is phosphorylated at S143 in the SH-SY5Y cells, which affected its interaction with the proteasomes. Expression of S143D-USP14 mutant or wildtype-USP14 reduced the levels of pS129 α-syn, α-syn and ROS in the USP14 deficient cells. These results demonstrate that USP14 plays a role in the clearance of pS129 α-syn, α-syn, and in reducing ROS in human SH-SY5Y dopaminergic cells that can be of importance in PD.

## Results

We deleted *USP14* in human SH-SY5Y dopaminergic cells by using CRISPR/Cas9 to target exon 2 of *USP14* gene and cell clones obtained showed no expression of USP14 (**Supplementary Figure 1A**).

### The 26S proteasome and 20S CP chymotrypsin-like activity are reduced in USP14-deleted cells

Proteasome 19S RP and 20S CP subcomplexes assemble to form the mature single or double-capped 26S/30S proteasome that perform ATP-and ubiquitin-dependent degradation of protein substrates. The ubiquitinated proteins upon binding to USP14 can accelerate proteasome-dependent degradation ^14, 21^. We cultured control and USP14 lacking cells to study the activity of the proteasome in cell lysates, separately for the 26S/30S proteasome holoenzymes and free 20S CP (see Methods). Data showed that the 26S/30S proteasome activity was downregulated in USP14-deleted cells compared with controls, most prominent in cells cultured for 48h (**Figure 1A and Supplementary Figure 1B**). Examining the free 20S CP chymotrypsin-like peptidase activity in the presence or absence of 0.02% sodium-dodecyl sulfate (SDS) to relieve the 20S CP gating ^36^ revealed a reduction in basal 20S CP activity in USP14 lacking cells, which was abrogated by SDS (**Figure 1A and Supplementary Figure 1B**). Analysis with native PAGE immunoblotting (IB) showed a reduction in the 26S proteasome complex, whilst the free 20S CP complex was increased in USP14-lacking cells (**Figure 1B and Supplementary Figure 1C**).

**Figure 1.**
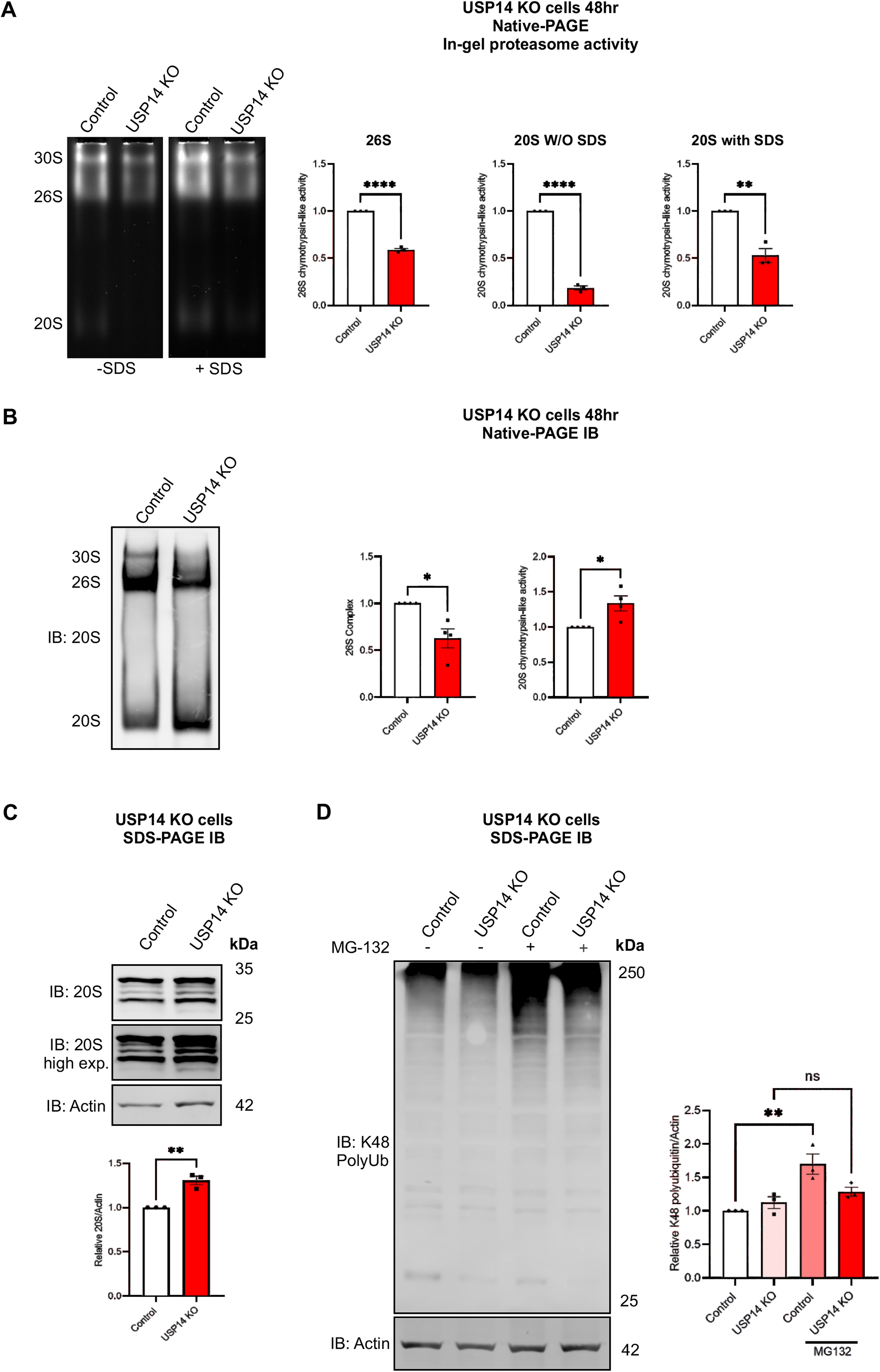
The 26S Proteasome activity and related protein complexes are reduced in USP14-deleted cells. Control SH-SY5Y and USP14-deleted cells were cultured for 48h as described in Methods (**A, B, C**). They were then treated for 5h with 20µM MG132, or DMSO as negative control, and analyzed further as indicated below. **(A)** Native-gel electrophoresis followed by in-gel activity assay was done as described in Methods for 26S/30S proteasome and 20S CP chymotrypsin-like activity. Left panels, UV-exposed native gel in the presence or absence of SDS to detect the activity of free 20S CP. Right panels, quantification of the densitometry ratio of 26S/30S proteasome, 20S CP activity without (W/O) or with SDS in USP14-deleted cells normalized to controls. Values are means ± S.E.M. **p ≤ 0.01, ***p ≤ 0.001, ****p ≤ 0.0001, n=3. **(B)** Native-gel electrophoresis followed by immunoblotting using a 20S antibody cocktail. Right panels, quantification of the densitometry ratio of 26S/30S and 20S complex in USP14-deleted cells normalized to controls. Values are means ± S.E.M. *p ≤ 0.05. n=4. **(C)** Immunoblotting using the antibody for 20S CP subunit proteins. Right panel, quantification of five 20S CP protein subunits normalized to β-actin. **(D)** Immunoblotting for K48-linked polyubiquitin chains antibody. Right panel, quantification of the K48-polyubiquitin smear normalized to β-actin. **(C-D)** Values are means ± S.E.M. **p ≤ 0.01, n=3.

Immunoblotting with SDS-PAGE using an 20S antibody cocktail revealed increases in the 20S CP subunits α5/α7, β1, β5, and β7 in USP14 lacking cells (**Figure 1C**). Notably, there were no changes in the 19S RP USP14 binding proteins, PSMD2 and PSMC2, or in UCHL5, a proteasome-associated DUB (**Supplementary Figure 1D and 1E**).

Protein substrates targeted for proteasome degradation often carry Lysine-48 (K48) linked ubiquitin chains ^37^. Inhibiting the proteasomes using MG132 revealed K48-linked polyubiquitinated proteins accumulation, shown as a smear, and this was significantly reduced in USP14-deleted cells (**Figure 1D**).

### Autophagy flux is increased in USP14-deleted cells

The proteasome and autophagy network are tightly interconnected ^38^, with USP14 playing a role in this crosstalk ^39, 40, 41^. Analysis of autophagy flux using chloroquine diphosphate (CQ) to inhibit the fusion of lysosomes with autophagosomes revealed increases in the lipidated ATG8 proteins, GABARAP-II (**Figure 2A**) and LC3B-II (**Figure 2B**) in cells lacking USP14. This indicated that the autophagy flux is elevated in USP14-deleted cells compared with controls.

**Figure 2.**
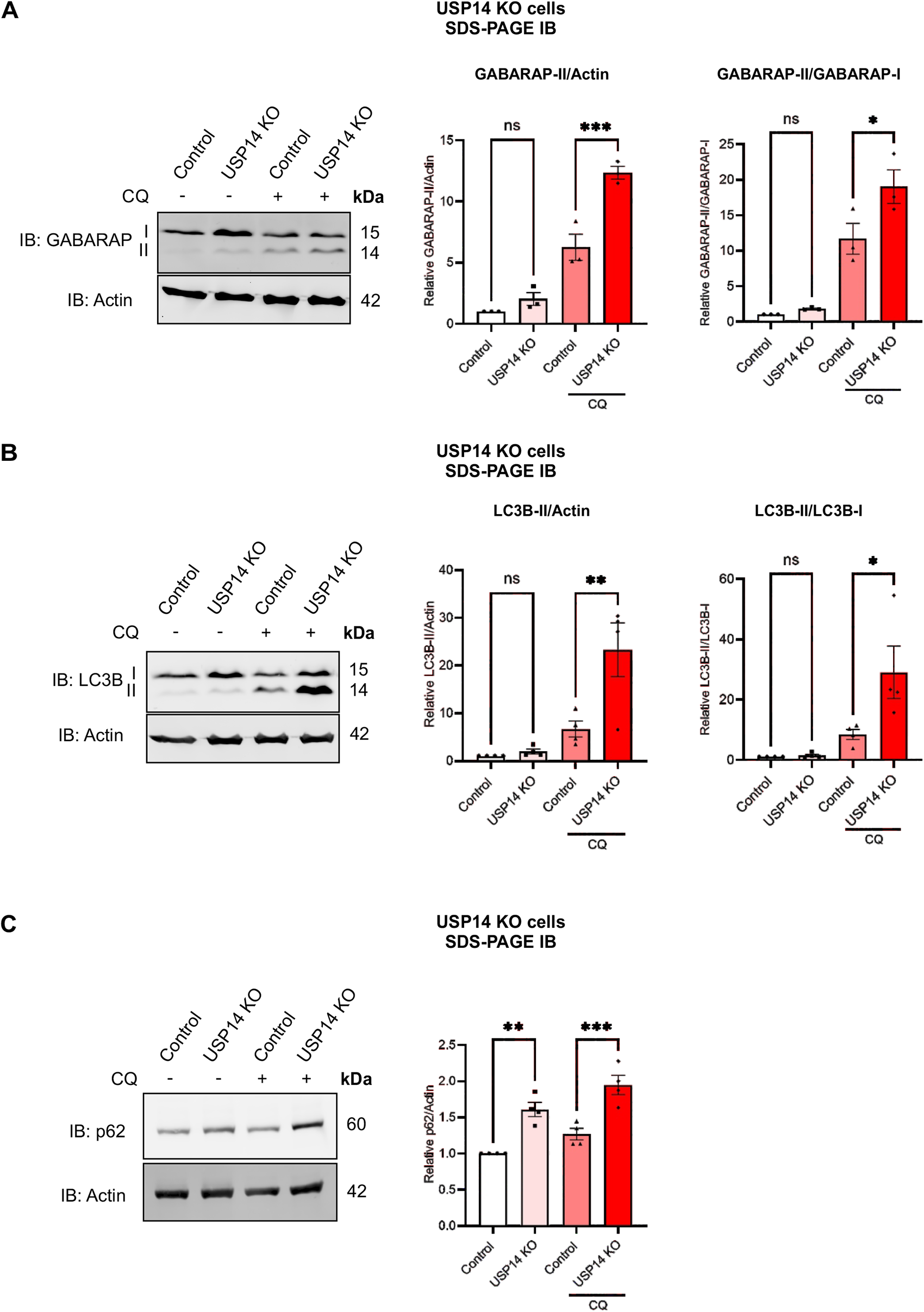
Increase in autophagy flux in USP14-deleted cells. Control and USP14-deleted cells were treated with 200µM chloroquine diphosphate (CQ) for 4h to inhibit autophagic flux and analyzed by immunoblotting. **(A)** Left panel, immunoblot of GABARAP. Right panels, quantification of the densitometry ratio of GABARAP-II normalized to β-actin or GABARAP-I. **(B)** Left panel, immunoblot of LC3B. Right panels, quantification of the densitometry ratio of LC3B-II normalized to β-actin or LC3B-I. **(C)** Left panel, immunoblot of p62. Right panel, quantification of the densitometry ratio of p62 normalized to β-actin. Values are means ± S.E.M. **p ≤ 0.01, ***-p ≤ 0.001, n=3-4.

Interestingly we observed that the autophagy receptor SQSTM1/p62 ^42^ was increased in USP14-deleted cells compared with controls (**Figure 2C**), and inhibiting the autophagy flux led to further elevation in p62. These results suggest a complex regulation of p62 in the USP14-deleted cells likely due to changes in both autophagy and proteasomes.

### USP14-deleted cells show elevated TFEB and CLEAR signaling

A central action of USP14 is to regulate the ubiquitination status of targets proteins to affect their degradation and intra-cellular signaling. To screen for proteins and pathways that are affected by *USP14* deletion, we performed LC-MS based proteome profiling with control and USP14-deleted cells lysates. Differentially Expressed Proteins (DEPs) with statistically significant changes in expression were further examined by the Ingenuity Pathways Analysis (IPA) (**Figure 3A-B**). This revealed overall, 356 different canonical pathways that were enriched or downregulated in USP14-deleted cells compared with controls (**Supplementary File 1 S1**). In addition to 26S proteasome degradation, ubiquitin metabolism, and chaperones, HSP90 family (association score = 32) (**Figure 3A**), top cellular pathways altered included the Coordinated Lysosomal Expression and Regulation (CLEAR) signaling (predicted to be activated; positive z-score = 3.162) (**Figure 3B**). The IPA analysis predicted the activation of the transcription factor EB (TFEB) in USP14-lacking cells, along with an elevation in 11 lysosomal enzymes (**Supplementary Figure 2**). TFEB is a master regulator of lysosomal biogenesis and functions ^43, 44, 45^, and immunoblotting confirmed an increase in TFEB in USP14 deficient cells compared with controls (**Figure 3C**). In line with this, the lysosomal enzymes, Glucocerebrosidase-1 (GBA1) and Cathepsin D were also elevated in USP14 lacking cells (**Figure 3D**). These results reveal a novel crosstalk between proteasome dysfunction and enhanced lysosomal activities via TFEB that may have functional significance for proteostasis.

**Figure 3.**
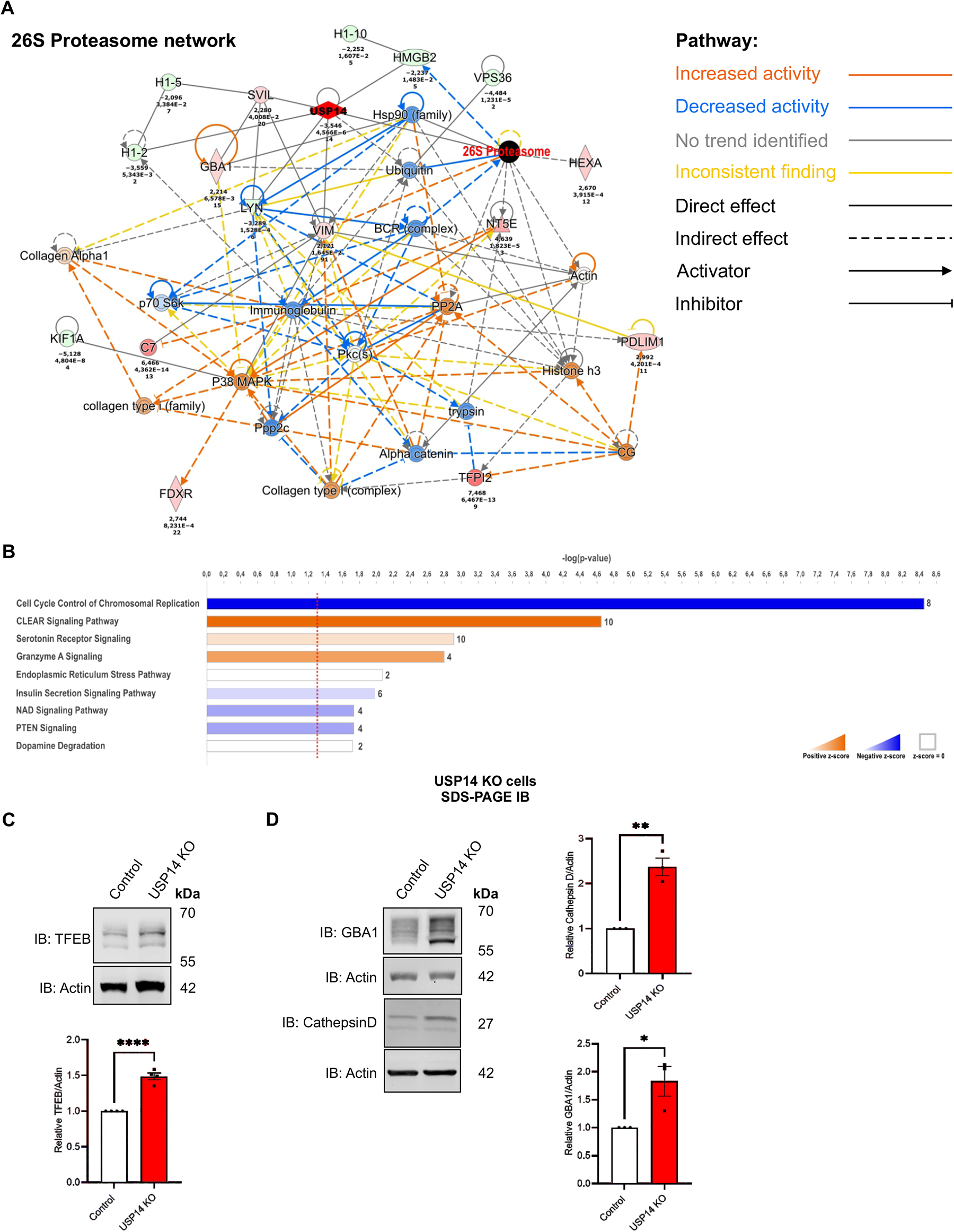
Cellular proteins and signaling networks are altered in USP14-deleted cells. Proteomic analyses of control and USP14-lacking cells were done as described in Methods **(A)** Network of interacting proteins related to the proteasome. The USP14 associated node is highlighted by red diamond and the 26S proteasome node by black circle. The color-coding describing changes in activity (red, blue) and direct of interaction is given on the right. The values below distinct proteins refer to changes in expression between USP14 deleted and control cells. **(B)** Top canonical pathways identified have been plotted as histograms with the number of proteins in each pathway shown to the right. The data was generated using the IPA software from the list of differentially expressed proteins (DEPs) between USP14ablated and control cells using n=4 biological repeats. The entire list of DEPs is provided in the Supplementary data file 1. The plot is represented as a fold change in p-value on the Y-axis and a threshold of p-value ≥ 1.3 was considered significant. DEP-Differentially expressed protein, IPA-Ingenuity pathway analysis **(C-D)** Cell lysates from control and USP14-deleted cells were subjected to immunoblotting as shown below, **(C)** Upper panel, immunoblot TFEB. Lower panel, quantification of the densitometry ratio of TFEB normalized to β-actin. Values are means ± S.E.M. ****p ≤ 0.0001, n=4. **(D)** Left panel, immunoblots GBA1 and Cathepsin D. Right panels, quantification of the densitometry ratio of GBA1 and Cathepsin D normalized to their respective β-actin. Values are means ± S.E.M. *p ≤ 0.05, **p ≤ 0.01, n=3.

### Mitochondria are elongated in USP14 deficient cells with increases in ROS

Studying mitochondria using MitoTracker in the SH-SY5Y dopaminergic cells, we observed an increase in mitochondrial branch length in USP14-deleted cells compared with controls (**Figure 4A**). In accordance with this, transmission electron microscopy (TEM) showed more elongated mitochondrial profiles in USP14-deficient cells (**Figure 4B**). The USP14-deficient cells also displayed elevated dark electron-dense vesicular structures (red asterisk) that likely represent lysosomes/autophagosomes accumulating in these cells (**Figure 4B**).

**Figure 4.**
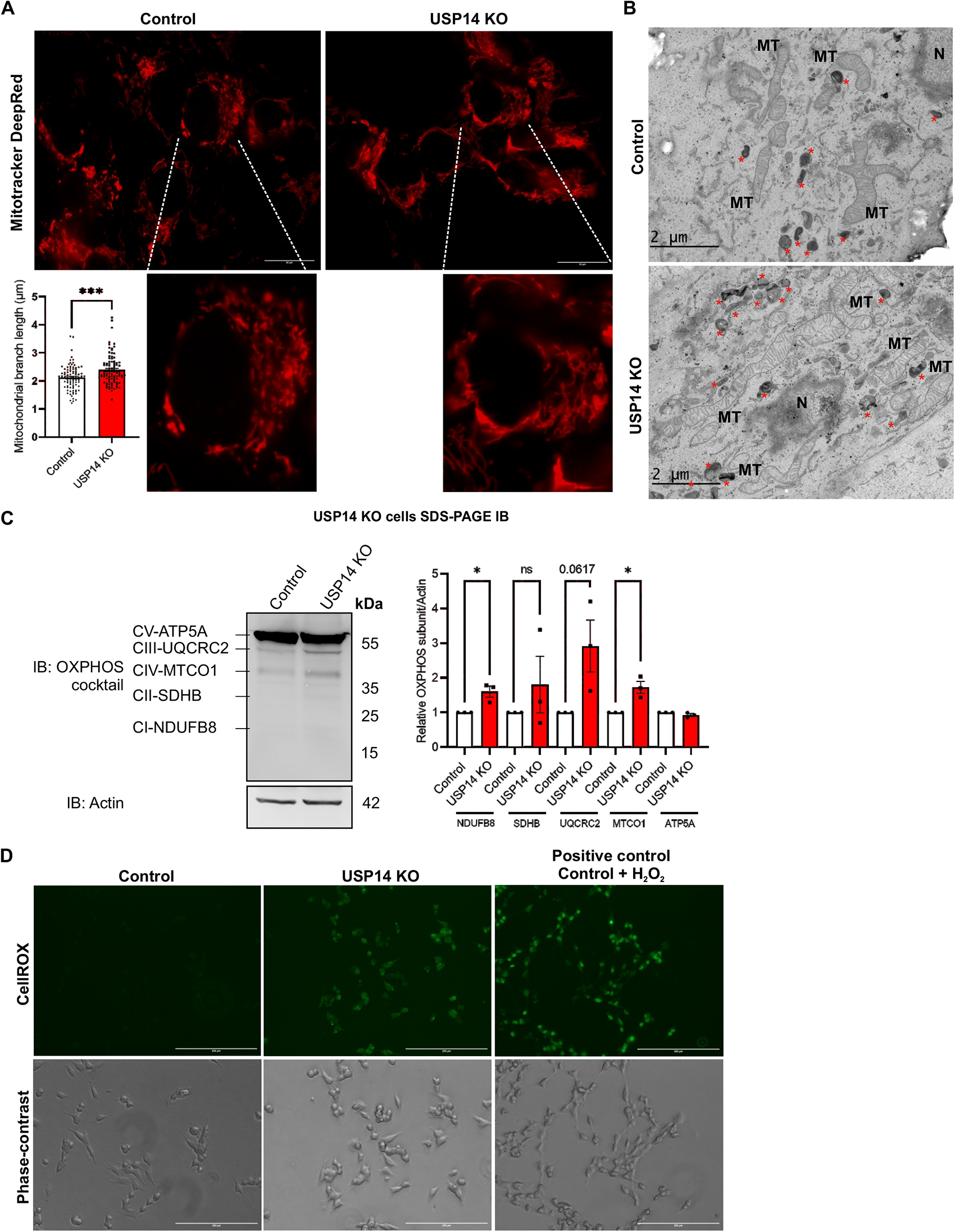
Elongation of mitochondria and elevated reactive oxygen species in USP14 deficient cells Control and USP14-deleted cells were cultured and analyzed as indicated below. **(A)** Live-cell imaging using Mitotracker DeepRed FM and a 100x objective of Nikon-Eclipse Ti-E inverted wide-field microscope equipped with an environmental chamber. Bottom panels, higher magnification. Left-bottom panel, quantification of mitochondrial branch length in µM from 90-100 control and USP14 deleted cells. Scale bar: 20µM. Typical experiment is shown and was repeated three times with similar results. **(B)** EM imaging was done as described in Methods. Note elongated mitochondria (MT) in USP14 deleted cells. Red stars * mark the presence of electron-dense lysosomes/ autophagosomes/ vesicles that were increased in the USP14-deleted cells compared with controls. N represents nuclear compartment. The experiment was repeated with similar results. **(C)** Left panel, immunoblot using an OXPHOS antibody cocktail. Right panel, quantification of the densitometry ratio of CI-CV subunits (CI-NDUFB8, CII-SDHB, CIII-UQCRC2, CIV-MTCO1 and CV-ATP5A) normalized to β-actin. Values are means ± S.E.M. *p ≤ 0.05, ns= not significant. n=3. **(D)** Live-cell imaging using CellROX Green and a 20x objective of EVOS FL microscope to visualize reactive oxygen species (ROS). Note increases of ROS in USP14-deleted cells Control cells treated with 5µM H_2_O_2_ for 90 min served as a positive control for oxidative stress. Top, CellROX Green. Bottom, phase-contrast images. Scale bar: 200µM. Typical experiment is shown and was repeated three times with similar results.

Immunoblotting for oxidative phosphorylation (OXPHOS) subunits using an antibody cocktail revealed increases in subunits of complex 1 (CI, NDUFB8), complex 3 (CIII, UQCRC2), and complex 4 (CIV, MTCO1) with no changes in complex 2 (CII, SDHB) or complex 5 (CV, ATP5A) (**Figure 4C**). Analysis of the basal mitochondrial oxygen consumption rate using the Seahorse^TM^ assay (Agilent Technology USA) revealed no clear differences between control and USP14-deficient cells (**Supplementary Figure 3A**).

Mitochondria are important sites to produce intracellular reactive oxygen species (ROS) as a byproduct of oxidative phosphorylation mainly at mitochondrial complex 3 ^46, 47^. Live-cell imaging using CellROX staining revealed increased ROS levels in USP14-deleted cells compared with controls (**Figure 4D**). As a positive control, we added H_2_O_2_ which elevated ROS in the control cells (**Figure 4D**).

### Total α-synuclein and pS129-α-syn are increased in USP14 deficient cells; the role of proteasome and oxidative stress

PD is associated with α-syn accumulation in neurons by mechanisms that are not fully understood. Utilizing immunoblotting, we observed that α-syn (**Figure 5A**) and PD pathology-associated pS129 α-syn (**Figure 5B**) levels are increased in SH-SY5Y cells lacking USP14 compared with controls. To investigate which pathways are involved in the degradation of α-syn, we used inhibitors for the proteasome (MG132), or autophagy (CQ). The accumulation of pS129-α-syn in response to proteasome inhibition was significantly higher in the USP14-deleted cells compared to controls (**Figure 5C**). Inhibiting autophagy increased α-syn only in the control cells, while USP14-deleted cells did not show any increase (**Figure 5D**). These results demonstrate that α-syn and pS129 α-syn accumulate in SH-SY5Y cells lacking USP14, correlating with defective proteasome degradation at basal levels. Notably, inhibiting the proteasomes by MG132 led to higher increases in pS129 α-syn in cells lacking USP14.

**Figure 5.**
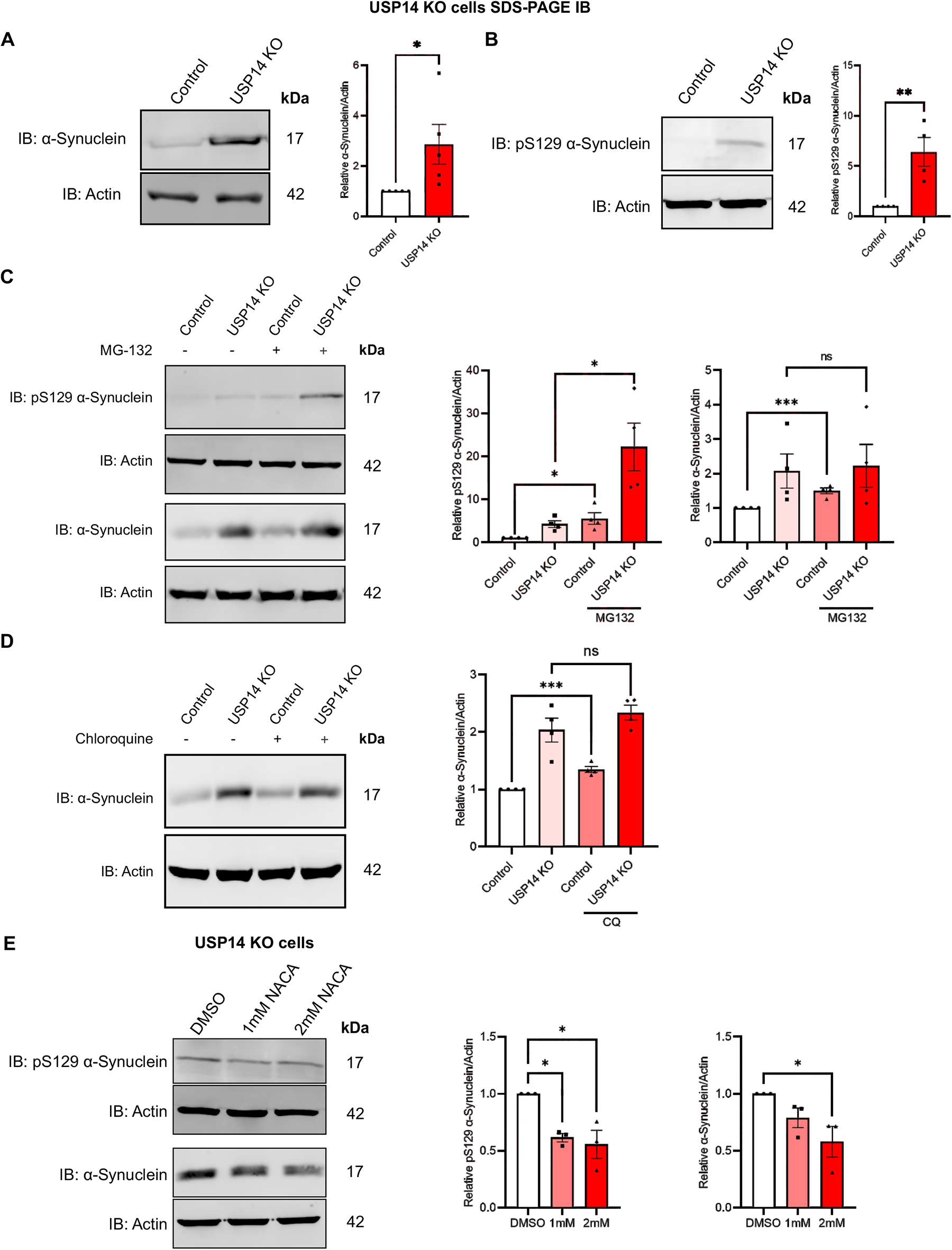
Total α-synuclein and pS129-α-syn levels are increased in USP14-deleted cells Control and USP14-deleted cells were analyzed as described below. **(A-B)** Left panels, immunoblots. Note increases in α-syn (A) and pS129 α-syn (B) levels in USP14-deleted cells. Right panels, quantification of the densitometry ratio of α-syn and pS129 α-syn normalized to their respective β-actin. Values are means ± S.E.M. *p ≤ 0.05, **p ≤ 0.01, n=4-5. **(C)** Treatment of cells with 20µM MG132 for 4h. Left panels, immunoblots of pS129 α-syn and α-syn. Right panels, quantification of the densitometry ratio of pS129 α-syn and α-syn normalized to their respective β-actin. Values are means ± S.E.M. *p ≤ 0.05, ***p ≤ 0.001, n=4. **(D)** Treatment of cells with 200µM Chloroquine (CQ) for 4h. Left panel, immunoblots of α-syn. Right panel, quantification of the densitometry ratio of α-syn normalized to β-actin. Values are means ± S.E.M. ***-p ≤ 0.001, n=4 **(E)** Treatment of USP14-deleted cells with 1 or 2mM N-acetylcysteine-amide (NACA) for 20h. Left panel, immunoblots. Right panels, quantification of the densitometry ratio of pS129 α-syn and α-syn normalized to their respective β-actin. Values are means ± S.E.M. *p ≤ 0.05, n=3.

To study whether oxidative stress contributes to the α-syn levels in the USP14-deficient cells, we utilized N-acetylcysteine amide (NACA) which reduced pS129 α-syn and α-syn levels in the USP14-deficient cells (**Figure 5E**) concomitant with decreasing ROS (**Supplementary Figure 3C**).

### USP14 is phosphorylated at S143 affecting its structure and functions

α-Syn levels and oxidative stress were increased in USP14 KO cells indicating a role of USP14 in the viability of SH-SY5Y dopaminergic. Post-translational modification (PTM) is one way to influence protein functions and interactions. To examine PTMs in USP14, we performed a quantitative phospho-proteomic study using overexpression of Flag-WT-USP14 in SH-SY5Y cells, followed by affinity enrichment, and analysis using LC-MS/MS (**Figure 6A**). Proteome Discoverer 2.5 (Thermo Scientific) identified putative phosphorylation sites in USP14 at several residues of which S143 phosphorylation was reliably recurring. Sequence alignment showed that S143 residue in USP14 sequence is conserved across eukaryotes including the yeast homolog, UBP6 (**Figure 6B**). To give insights into the structure of USP14, we performed molecular dynamics (MD) simulations of USP14 phosphorylated at the S143 (pS143-USP14) and USP14 mutated at the S143 residue using the Maestro 13.0 InterfaceDesmond package of the Schrödinger LLC. MD simulations with the 30ns length were carried out for each structure. Protein secondary structure comparisons of WT-USP14 with pS143-USP14 (**Figure 6C**), and WT-USP14 with S143A-USP14, S143D-USP14 (**Supplementary Figure 4A**) suggested that the S143 phosphorylation affects the UBL domain conformations in USP14. An additional run performed with 50ns length confirmed the stability of the MD simulations and the results obtained (data not shown).

**Figure 6.**
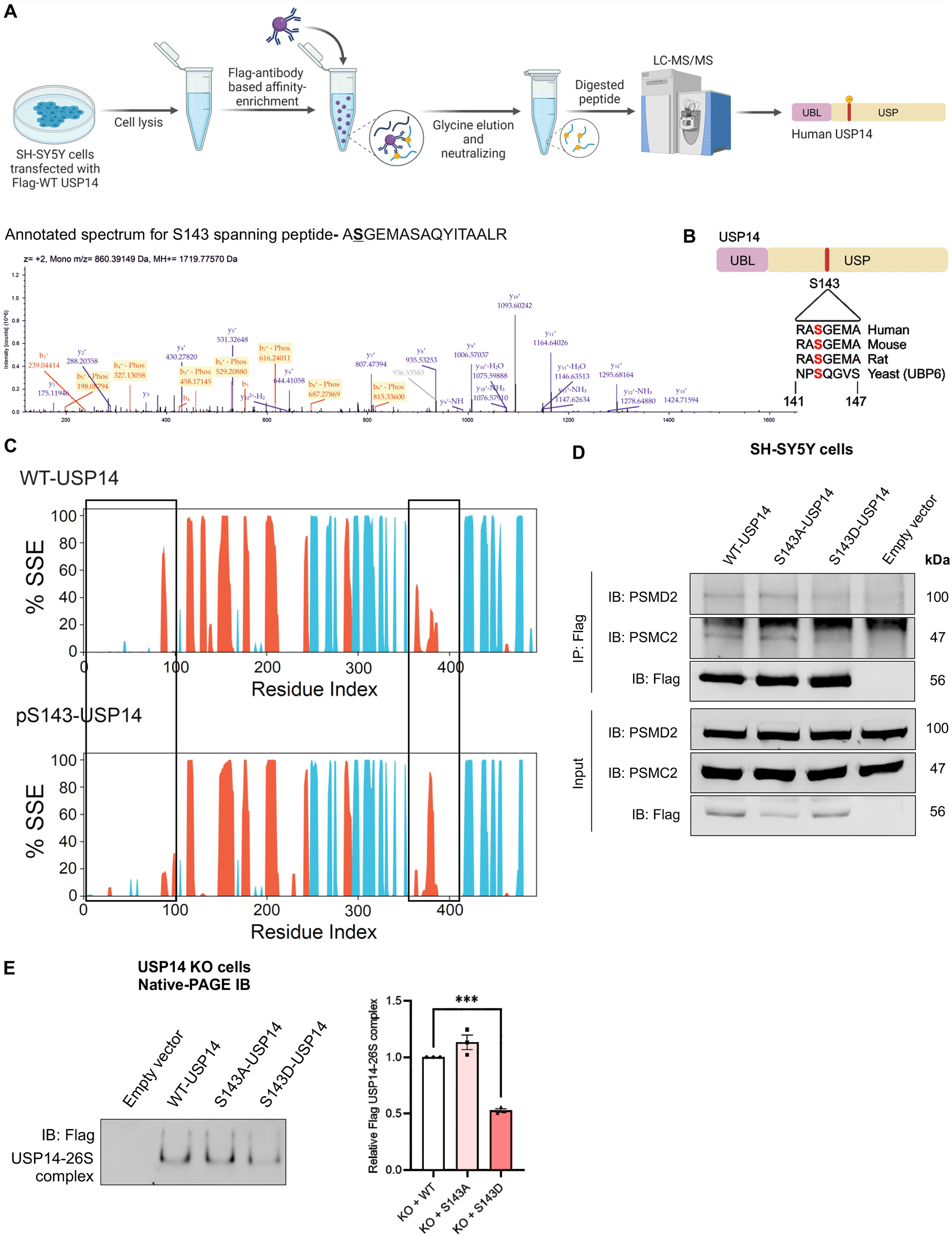
USP14 is phosphorylated at S143 as identified by quantitative mass spectrometry. **(A)** Workflow for phospho-proteomic experiments. SH-SY5Y cells overexpressing Flag-WT-USP14 were lysed and processed as described in Methods. Immunoprecipitated Flag-WT-USP14 was eluted using glycine, neutralized, enzymatically digested, and processed for LC-MS/MS analysis of phospho-peptides. Below panel, LC-MS/MS annotated mass spectrum of A**<u>S</u>**GEMASAQYITAALR peptide spanning pS143 residue identified from a representative experiment. The experiment was repeated four times with similar results. **(B)** Graphical illustration of human USP14 showing the N-terminal ubiquitin-like (UBL) and C-terminal ubiquitin-specific protease (USP) domains. Lower half depicts the sequence alignment and the conservation of the S143 residue across eukaryotes. **(C)** Protein secondary structure elements (SSE) distribution by residue index throughout the protein structure monitored throughout the MD simulation are shown for WT-USP14 and USP14 phosphorylated at S143 (pS143-USP14). α-helices and β-strands are shown in orange and blue, respectively. The black rectangular boxes indicate the areas with the largest conformational changes. MD simulations were done as described in Methods. **(D)** SH-SY5Y cells were transfected with Flag-tagged WT-USP14, S143A-USP14 or S143D-USP14 for 24hr, and lysates immunoprecipitated with Flag antibody. The eluates were analyzed by immunoblotting using anti-PSMD2, anti-PSMC2, and anti-Flag antibodies. Input lanes, total cell lysate. A representative experiment is shown and was repeated three times with similar results. Note: Reduced interaction of Flag S143D-USP14 with the 19S RP components PSMD2 and PSMC2 **(E)** USP14-deleted were reconstituted with Flag-tagged WT-USP14 or S143A-USP14 or S143D-USP14 for 24hr and lysates were analyzed by native-gel electrophoresis followed by immunoblotting for Flag antibody to detect Flag-USP14 bound 26S proteasomes. Right panel shows quantification of the densitometry ratio of Flag-S143A-USP14 or Flag-S143D-USP14 containing 26S complexes normalized to Flag-WT-USP14 containing 26S complexes. Values are means ± S.E.M. ****p ≤ 0.0001, n=3. Note: Reduction in Flag-tagged S143D-USP14 bound to 26S proteasomes.

We next generated phospho-deficient (S to A) S143A-USP14 and phospho-mimetic (S to D) S143D-USP14 mutants to perform biochemical and functional validations. Immunoprecipitation using wildtype USP14 and S143 mutants revealed a reduced binding of the S143D-USP14 mutant to the proteins, PSMD2 and PSMC2 (**Figure 6D**). Native-PAGE immunoblotting of lysates from USP14-deleted cells transfected with Flag-tagged WT-USP14, or S143-USP14 mutants revealed a lower interaction of the S143D-USP14 with the 26S proteasome complexes (**Figure 6E**). Together these results demonstrate a reduced binding of the S143D-USP14 mutant to the proteasome in line with the prediction form the MD.

To investigate whether the phosphorylation at S143 may influence the catalytic activity of USP14, we performed a DUB activity assay utilizing UbVME as a substrate in parental SH-SY5Y cells expressing either Flag-tagged WT-USP14, or USP14 S143 mutants. Immunoblotting for UbVME-bound USP14 revealed that S143D-USP14 had a slightly higher activity compared to WT-USP14 and S143A-USP14 **(Supplementary Figure 4B).** However, it should be remembered that UbVME is a synthetic substrate and the behavior of the USP14 towards natural substrate may be different.

### USP14 re-expression reduces ROS, pS129 α-synuclein, and α-synuclein USP14-deleted cells

To elucidate the functional role of S143 phosphorylation further, we reconstituted WT-USP14, S143A-USP14 or S143D-USP14 in the USP14-deleted cells. Live-cell imaging of CellROX staining showed that the WT-USP14 and the S143D-USP14 mutant reduces ROS levels in the USP14-deleted cells while the S143-USP14 had no effect (**Figure 7A**). Similarly, WT-USP14 and pS143-S143D reconstitution into USP14-deleted cells also reduced the levels of pS129 α-syn (**Figure 7B**) and α-syn (**Figure 7C**), while the S143A-USP14 mutant had no statistically significant effect.

**Figure 7.**
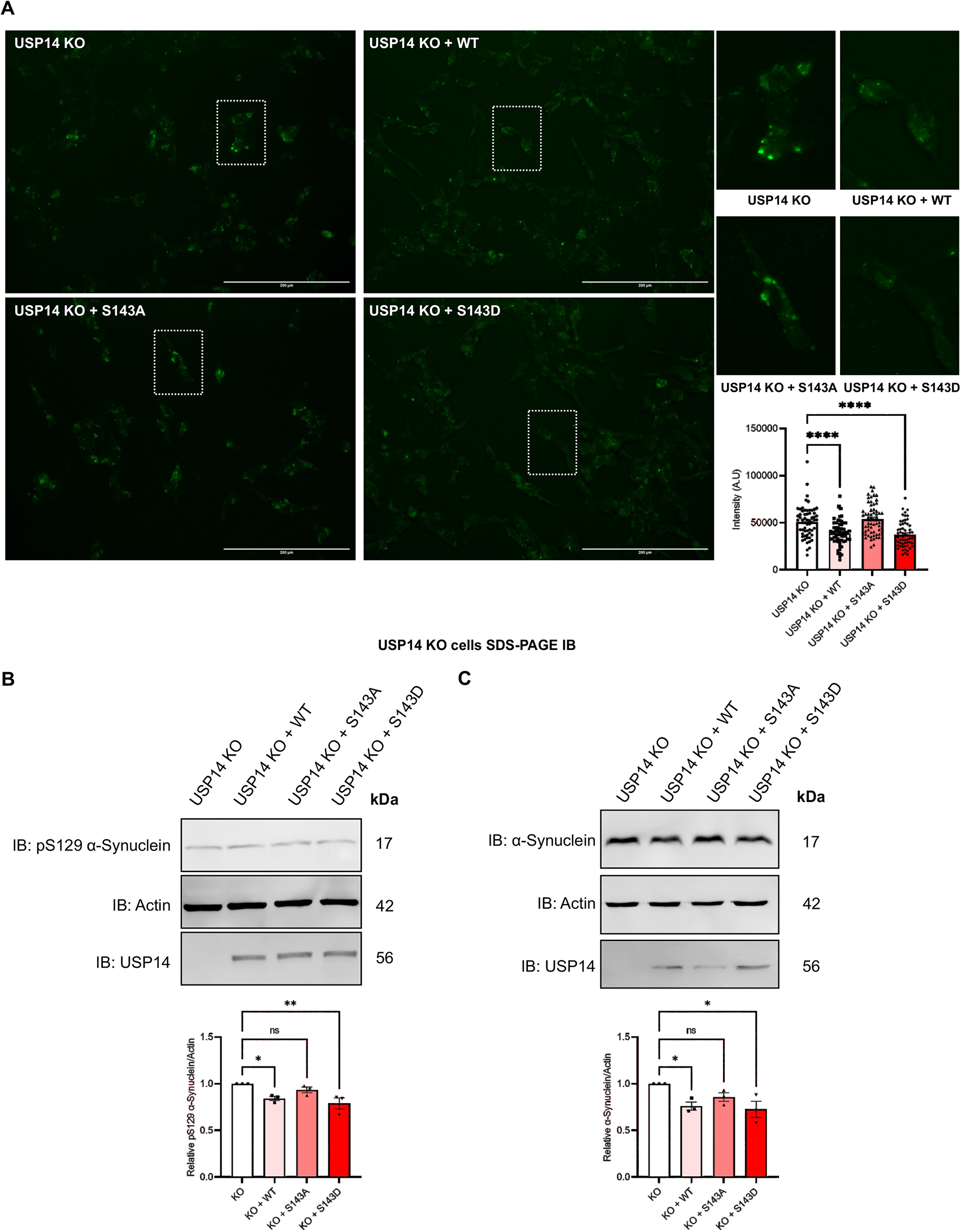
USP14 expression reduces ROS and α-syn levels in USP14-deleted cells. **(A-C)** Reconstitution experiments. USP14-deleted cells were transfected with plasmids expressing USP14, Flag-tagged wildtype (WT)-USP14 or the mutant USP14, S143A-USP14, and S143D-USP14. 24h later, cells were analyzed as below. **(A)** Live-cell imaging of CellROX Green stain. Scale bar: 200µM. Right panels show representative cells (marked by dotted white rectangle). Note: a reduction in ROS using WT-USP14 and S143D-USP14 mutants but not with the S143A-USP14 mutant. Right panel histogram. Quantification was done by analyzing the intensity of 50-60 cells in each well and plotted as a histogram. A typical experiment is shown and repeated with similar results. Values are means ± S.E.M. ****p ≤ 0.0001. **(B)** Top, immunoblots. Bottom, quantification of the densitometry ratio of pS129 α-syn normalized to β-actin. Note: pS129 α-syn level is reduced by reconstitution of WT-USP14 and S143D-USP14 in USP14-deleted cells. Values are means ± S.E.M. *p ≤ 0.05, **p ≤ 0.01, ns-not significant, n=3. **(C)** Top, immunoblots. Bottom, quantification of the densitometry ratio of α-syn normalized to β-actin. Note: α-syn level is reduced by reconstitution of WT-USP14 and S143D-USP14 in USP14-deleted cells. Values are means ± S.E.M. *p ≤ 0.05, ns-not significant, n=3.

## Discussion

In the present work, we provide evidence that USP14 is a critical regulator of α-synuclein (α-syn) clearance and oxidative stress in SH-SY5Y dopaminergic cells. At the molecular level, deleting USP14 inhibited the chymotrypsin-like activity of the 26S proteasome, and 20S CP while activating the TFEB-mediated CLEAR/autophagy signaling pathway. USP14-ablated cells exhibited an increase in oxidative stress, which could negatively impact 26S proteasome activity and aggravate mitochondrial dysfunctions. Clearance of α-syn and pS129 α-syn was dependent on proteasome activity and on reducing ROS in these SH-SY5Y dopaminergic cells. Furthermore, USP14 is phosphorylated at S143, affecting its interaction with 26S Proteasomes, and is important for counteracting ROS, pS129 α-syn, and α-syn levels in USP14-ablated cells.

USP14 is important for regulating the 26S proteasome function ^11, 17, 18^. In yeast, UBP6 (USP14 homolog) induces the 19S RP ATPase ring activation, ATP hydrolysis, and thus, activating 26S proteasome peptidase activity ^21^. On the other hand, purified 26S Proteasomes from USP14-deleted MEF cells showed higher activity for all three peptidases and ATP-hydrolysis ^13^. This indicates that the functional interactions of USP14 with the proteasome is complex, and context-dependent based on the availability of ubiquitinated substrates and other factors ^18^ .

In our study, USP14-deleted SH-SY5Y neuronal cells exhibited a loss in 26S proteasome and basal 20S CP activity, and increased 20S CP gating. Correspondingly, the amount of the individual 20S CP subunits was elevated, which could translate to increased gating of the free 20S CP complex and a related decrease in the 20S CP basal activity in USP14-deleted cells. In addition, we did not observe compensatory changes in UCHL5 DUB that regulates 26S proteasomes, or in the levels of USP14 interacting proteins in 19S RP, PSMD2, or PSMC2. Together these findings support a critical role of USP14 in activating the 26S Proteasomes and the free 20S CP in SH-SY5Y neuronal cells. This is in line with the findings in yeast that UBP6/USP14 is important for 26S proteasome activation ^21^. It would be important to study the involvement of shuttle factors such as UBQLN2 ^48^ in USP14-dependent regulation of 26S Proteasomes in different cell types and in the clearance of aggregation-prone proteins, such as α-Synuclein/ pS129 α-Synuclein.

TFEB is a master regulator of autophagy-related proteins and lysosomal biogenesis encompassing the previously identified CLEAR network ^43, 44, 49^. TFEB itself is a substrate for the proteasome and TFEB phosphorylation controls its nuclear translocation and transcription factor activity ^40^. E3 ligase CHIP/STUB1 regulates TFEB while less is known about the role of DUBs in TFEB regulation ^50^. Global proteome profiling identified TFEB-mediated signaling to be increased in USP14-deleted cells. TFEB is elevated in USP14-deleted cells together with the downstream ATG8 proteins, and the lysosomal enzymes, β-Glucocerebrosidase and cathepsin-D.

Biochemical validations and EM imaging corroborated these findings showing elevated lipidation of LC3B and GABARAP, accumulating lysosomes, and autophagic flux in dopaminergic cells lacking USP14 probably as a compensation to proteasome inhibition. The elevated TFEB and autophagy in conjunction with higher levels of α-syn and pS129 α-syn were unexpected but in line with the clearance of these molecules via proteasomes. This is also at slight variance with study showing the benefit of overexpressing TFEB in an *in-vivo* model of α-syn toxicity ^51^. However, a recent study reported an increase in the abundance of lysosomes in brain dopaminergic neurons after injection with α-syn pre-formed fibrils ^52^. This resembles the situation in the SH-SY5Y dopaminergic cells devoid of USP14. However, more data is warranted to resolve the roles of lysosomes and autophagy pathways related to α-syn in dopaminergic neurons and in models of PD.

Mechanistically, USP14 can stabilize UVRAG levels ^39^ and reduce K63-linked polyubiquitin chains on Beclin-1 to regulate downstream autophagy ^53^. We have further shown that W58A-USP14 mutant, incapable of binding to the proteasome, increases GABARAP-positive structures related to selective autophagy in striatal neuronal cells ^25^. In addition, inhibiting USP14 with IU1 in thyroid cancer cells, augmented LC3B-lipidation, and the autophagy flux ^30^. The present results demonstrate that USP14 is important in coordinating responses between proteasomes and autophagy by regulating TFEB and the CLEAR signaling network in the SH-SY5Y dopaminergic cells.

USP14-deleted cells showed elongation of mitochondria in comparison to controls. The elongation of mitochondria could partly be due to changes in cellular proteins regulating the dynamics of mitochondrial fission and fusion or lack of autophagosome recruitment to mitochondria. Preliminary data showed changes in the fission-inducing protein Drp1 phosphorylation, possibly causing a mitochondria fission defect in the USP14-deleted cells (data not shown). On the functional level, respiratory chain complex 1 and 3 OXPHOS subunits were altered in SH-SY5Y cells devoid of USP14. However, the cells had a largely intact respiratory capacity. Cellular ROS was increased in USP14-lacking cells compared with controls as shown by live-cell imaging of CellROX dye. Excessive ROS arising from mitochondrial respiration can lead to oxidative stress in PD and Huntingtońs disease (HD) ^54, 55^. Oxidative stress can further impact the assembly/disassembly of proteasomes ^56^ and clearance of oxidative-damaged proteins ^57^. The precise targets for the action of USP14 on mitochondria related to oxidative stress - 26S proteasome crosstalk remains to be studied. α-Syn aggregates can disrupt mitochondrial functions and result in oxidative stress in midbrain dopaminergic neurons susceptible to degeneration in PD ^9, 52^. More focus has recently been on understanding pS129 α-syn and its role in PD. Increase in pS129 α-syn correlates with early-onset proteasome defects in dopaminergic neuronal PD models ^58^. pS129 α-syn has also been identified to be part of α-syn aggregates and is present in α-syn mutant forms such as A53T ^31^.

We identified here that α-syn and pS129 α-syn levels are increased in USP14-lacking SH-SY5Y dopaminergic cells mostly due to the proteasomal defect. Notably, the elevated pS129 α-syn levels following proteasome inhibition compared to α-syn indicates a difference in the clearance of these molecular species in the SH-SY5Y dopaminergic cells. This could be due to the higher binding avidity for pS129 α-syn to proteasome or involve other proteins preferentially interacting with pS129 α-syn. α-Syn can undergo ubiquitin-independent degradation via the 20S CP owing to its intrinsically disordered region that is a determinant of 20S CP degradation ^59^. We observed a defect in 20S CP activity in the USP14-ablated cells that may contribute to the increases noted in α-syn and pS129 α-syn levels.

Oxidative stress is a major component of neurodegenerative diseases including PD. We noted that deleting USP14 in SH-SY5Y dopaminergic cells increases oxidative stress with elongated mitochondria. Addition of NACA as a synthetic antioxidant, decreased ROS, pS129 α-syn, and α-syn levels in the USP14-deleted cells. In addition, reconstituting WT-USP14 and phospho-mimetic S143D-USP14 mutant in USP14-ablated cells lowered α-syn and pS129 α-syn levels. Collectively these findings demonstrate that USP14, its phosphorylation at S143, and ROS are involved in the regulation of α-syn and pS129 α-syn clearance in SH-SY5Y dopaminergic cells.

Post-translational modifications are important determinants of protein structure and functions. USP14 can be phosphorylated at serine residue 432 (S432) in HEK293T cells downstream of the protein kinase Akt ^60^. We identify here that USP14 is phosphorylated at Serine 143 (S143) in SH-SY5Y dopaminergic cells, while S432 was not. This indicates that USP14 can be phosphorylated at different residues, which may vary between cell types and conditions. The catalytic groove pocket in USP14 comprises of: Cysteine 114, Histidine 435, and Aspartic acid 451 ^23^. The S143 phosphorylation identified in the present study lies within the region between the α2 and α3 helices of the USP14 catalytic domain (USP domain) ^23^. A key finding from our MD simulation was that S143 phosphorylation affects the UBL-domain conformation in USP14, which is important for the interaction with 26S Proteasomes ^61, 18, 22^.

Further investigation confirmed that the S143D-USP14 associated less with the 26S proteasome complex, and the 19S RP subunits PSMD2, and PSMC2. Recently, an NMR study reported a functional coupling of UBL and USP domains in the USP14 structure ^62^. In line with this, we observed that the S143 phosphorylation located within the USP domain impacts the UBL domain, regulating the interaction of USP14 with the 26S Proteasomes.

Importantly, reconstitution experiments elucidated that the S143D-USP14 mutant was able to reduce ROS levels in USP14-deleted cells, while the S143A-USP14 did not. Expression of WT-USP14 possibly upon S143 phosphorylation or other unidentified mechanisms could decrease ROS in USP14-lacking cells. Further studies are required to investigate the status of USP14 phosphorylation under different conditions by utilizing phospho-specific antibodies and the associated upstream kinases/phosphatases that regulate USP14 functions. Given the importance of USP14 in regulating key cellular functions, S143 phosphorylation could have a broader function in cell physiology. Elucidating factors involved in regulating USP14 and proteostasis may provide novel targets to interfere with protein aggregates in SH-SY5Y dopaminergic cells and PD.

## Materials and methods

### Cell culture and transfections

The SH-SY5Y human neuroblastoma cell line has been used as a culture model of dopaminergic neurons in studies of PD ^63^. SH-SY5Y cells were cultured in Dulbecco’s Modified Eagle Medium (DMEM) (Gibco, USA) supplemented with 10% fetal bovine serum (Gibco, USA), 7.5% NaHCO3, 100 mM NA-glutamine (Gibco, USA) and 100 mM penicillin-streptomycin (Gibco, USA) at 37°C in 5% CO2.

Cells were transfected with Linear Polyethylenimine 25.000 (PEI) (Polysciences, Inc., USA) or Fugene HD reagent (Promega, USA) following the manufacturer’s instructions. For PEI, stock PEI solution was prepared at a concentration of 1 mg/ml. For complexing the DNA with PEI or Fugene, they were combined at a ratio of 1:3 and added onto the cells. Following 24 - 48h of transfection, cells were further processed and used in experiments described below.

### CRISPR/Cas9 mediated *USP14* gene deletion

*USP14* gene deletion in SH-SY5Y cells was performed utilizing the Clustered regularly interspaced short palindromic repeats (CRISPR) Cas9 endonuclease (CRISPR-Cas9) system as previously described ^64^. Briefly, CRISPR design tool (https://crispr.mit.edu) was used to design guide RNAs (gRNAs) targeting the exon 2 of human *USP14* (common exon to most human USP14 transcripts). gRNA USP14_Forward: 5’ CACCGGTGAGCCTTGAATACCATTGG 3’ and gRNA

USP14_Reverse: 5’ AAACCCAATGGTATTCAAGGCTCACC 3’ were cloned into pSpCas9(BB)-2A-Puro vector (PX459, 62988, Addgene, USA) using FastDigest BpiI (Thermo Fisher Scientific, USA) and T4 DNA ligase (New England Biolabs, USA). The plasmids were sequenced and SH-SY5Y cells were cultured and transfected as described above. Control cells were transfected with pSpCas9(BB)-2A-Puro vector without USP14 specific gRNAs. At 48h post-transfection, puromycin-containing growth medium was added, and cells selected for using single-cell clones in 96-well plates to generate stable cell clones. Validation of the USP14 knock-out (KO) cell clones was conducted using immunoblotting with anti-USP14 antibody.

### Mutagenesis of USP14

Site-directed mutagenesis was done to generate Serine143 (S143) mutations in human *USP14* gene. at Genome Biology Unit (GBU), University of Helsinki. In brief, the USP14 entry clone from the human ORFeome collaboration library was used to perform mutagenesis and the constructs obtained were transferred into the 2Flag-pDEST-N (118371, Addgene, USA) vector using standard reaction protocol. The S143A-USP14 (loss-of-function) and S143D-USP14 (gain-of function) constructs were further sequenced and employed for cell cultures studies. The 2Flag-pDEST-N vector with no insert was utilized as a negative control for transfections.

### Molecular modeling of USP14

*Homology modeling of the structure of the USP14 and generation of USP14 mutant structures.* The 3D structure of the full-length USP14 was constructed using the hierarchical approach to protein structure and function prediction by I-TASSER (Iterative Threading ASSEmbly Refinement) server^65^. The structure with higher C-and TM-scores was selected for further modelling. The structures of the USP14 mutated at S143 residues (S143A and S143D) were generated using the Maestro 13.0 interface of the Schrödinger Software. The structure of the USP14 phosphorylated at S143 residue was generated using PyTMs plugin in The PyMOL Molecular Graphics System, Version 3.0 Schrödinger, LLC ^66^.

The 3D structure of the USP14 obtained by homology modelling was treated before further molecular dynamics (MD) simulation using Schrodinger Protein Preparation Wizard ^67^. Thereafter, the prepared structure of the protein was minimized using MD simulation by Desmond program package of Schrödinger LLC ^68^. The structure optimized by MD simulation was used for further modelling.

*Molecular dynamics (MD) simulations*. The MD simulations were carried out using the Desmond program package of Schrödinger LLC ^68, 69^. The MD simulations were performed similarly as described previously ^70^. The analysis of the MD trajectories was performed using the Simulation Interaction Diagram tool implemented in Desmond molecular dynamics package ^69^.

### Immunoblotting

Cells were washed with ice-cold PBS, twice and lysed in Radioimmunoprecipitation assay buffer (RIPA) buffer containing 150 mM NaCl, 1% Triton-X-100, 0.5% sodium deoxycholate, 1% SDS, 50 mM Tris-HCl, pH 7.4 supplemented with protease inhibitors (Roche, Switzerland) and phosphatase inhibitor (Phosphostop, Roche, Switzerland) ^25, 30^. Lysates were sonicated, centrifuged, and the supernatant protein concentration measured using BCA protein assay kit (Pierce, Thermo Fisher Scientific, USA). Equal amounts of proteins were separated on denaturing SDS-PAGE and blotted onto nitrocellulose membrane filters (Sartorius). The membranes were blocked and incubated with the primary antibodies overnight at 4°C with gentle agitation. The primary antibodies used included: USP14 (J6111-6D6, Sigma, Germany), α-synuclein (2642, CST, USA), pS129-α-synuclein (EP1536Y, Abcam, GB), 20S CP subunits cocktail (BML-PW8195, Enzo Lifesciences, USA), K48-linkage specific polyubiquitin (4289, CST), PSMD2 (sc-271775, SantaCruz, USA), PSMC2 (14395, CST), UCHL5 (sc-271002, SantaCruz, USA), TFEB (37785, CST, USA), GBA1 (sc-166407, SantaCruz, USA), GABARAP (13733, CST, USA), LC3B (3868, CST, USA), p62/SQSTM1 (P0067, Sigma, Germany), beta-Actin (A2066, Sigma, Germany), GAPDH (MAB374, Millipore, Germany), OXPHOS cocktail (MS604, Mitosciences, USA – Currently Abcam, GB)

Following incubation overnight, horse-radish peroxidase conjugated secondary antibody (1:2500, Jackson Immunoresearch Laboratories, USA) were added and the membranes incubated for 1h at room temperature with gentle agitation. Protein signals were detected using enhanced chemiluminescence substrate (Pierce, Thermo Fisher Scientific, USA). Immunoblots were quantified with ImageJ (NIH, Bethesda, USA) quantification software.

For immunoblotting of pS129-α-synuclein, the samples were heated and separated on BioRad 4-20% gradient gels and transferred onto nitro-cellulose membranes. Upon transfer, the membranes were fixed immediately in 0.4% PFA diluted in TBS. The membrane was then washed with TBS-T, blocked with 5% milk diluted in TBS-T and incubated in pS129-α-synuclein antibody for 1h, RT with gentle agitation. Following this, membranes were washed with TBS-T and incubated with HRP-conjugated anti-rabbit secondary antibodies for 1h, RT with gentle agitation. Post antibody incubation, membranes were washed with TBS-T and imaged using enhanced chemilumisescence substrate as above.

### Immunoprecipitation

Cells were cultured and lysed in IP-lysis buffer containing 50 mM Tris-HCl pH 7.7, 150 mM NaCl, 1% NP-40, and 0.5% sodium deoxycholate for 15min on ice followed by centrifugation at 10,000xg for 10min at 4°C ^25^. The supernatant was collected, and protein concentration was measured using BCA assay (Pierce, Thermo Fisher Scientific, USA).

The antibody-bead complexes were centrifuged, pelleted, and washed with IP-lysis buffer thrice. Immunoprecipitated proteins were eluted by the addition of 25µl 2x denaturing Laemmli buffer and heated to 98°C for 5 min. The eluates were then subjected to conventional denaturing SDS-PAGE and analyzed using immunoblotting for the indicated antibodies. The total cell lysates served as the input controls.

### Native-PAGE to perform in-gel proteasome activity assay and protein complex composition

20S CP and 26S/30S proteasome chymotrypsin-like activity, and proteasome complexes were analyzed using in-gel proteasome activity assay and immunoblotting followed by Native-PAGE, as described before ^30, 36, 71, 72^. Control and USP14-deleted SH-SY5Y cells received different treatments and lysed in Over Kleeft (OK) lysis buffer (50 mM Tris-HCl pH7.5, 2 mM DTT, 5 mM MgCl2, 10% glycerol, 2 mM ATP, and 0.05% Digitonin) by incubation on ice for 20min with intermittent vortexing. Lysates were then centrifuged at 27,670xg for 20min at 4°C, protein concentration were estimated as above, and an equal amount of protein was separated on Tris-Borate PAGE in a running buffer comprised of Tris-borate buffer supplemented with EDTA-Na2, ATP and MgCl2. Gels were run at a constant voltage of 120V approximately 2h 30min in the cold room. Following this, the gels were either transferred to a buffer containing in-gel activity substrate or buffer for performing immunoblotting.

For in-gel activity assay, gels were incubated for 15min, 37°C in the substrate buffer (50 mM Tris-HCl pH 7.4, 5 mM MgCl2, 1 mM ATP) supplemented with 100 mM of Suc-Leu-Leu-Val-Tyr-AMC (Bachem, Switzerland) substrate for chymotrypsin-like activity of 20S CP. Gel imaging was performed by exposing them to an excitation wavelength of 380 nm and emission wavelength of 460 nm. Upon imaging, the gels were incubated again for 15 min 37 °C in the above-mentioned substrate-containing buffer added with SDS to open the gating of free 20S CP ^36^. Furthermore, the gels were imaged again to obtain activated 20S CP activity in the presence of SDS.

For immunoblotting gels were incubated in conventional Tris-Glycine running buffer containing SDS for 10 min with gentle shaking and then transferred to the Tris-Glycine transfer buffer containing methanol. Standard protocol for wet-transfer was performed for 16h, 20V at 4°C followed by incubation with 20S (BML-PW8195, Enzo Lifesciences, USA), anti-USP14 antibody (J6111-6D6, Sigma, Germany).

### In-vivo DUB catalytic activity assessment using the Ub-VME substrate

The labelling assay for DUB activity was performed as described earlier ^12, 25^. SH-SY5Y cells were transfected with Flag-tagged WT-USP14 or S143A-USP14 or S143D-USP14. Cells were lysed after 24h using 50 mM Tris (pH 7.4), 250 mM sucrose, 5 mM MgCl2, 1mM DTT and 1mM ATP for 1h at 4°C, and 30 µg of cell lysates were incubated with 2 mM ubiquitin vinyl methyl ester (Enzo Life Science) for 1h at 37°C. The reaction was stopped by addition of Laemmli buffer and samples were analyzed by SDS-PAGE immunoblotting as above with Flag antibody to visualize the UbVME-bound Flag-tagged USP14.

### Sample preparation, protein digestion, and nano LC-MS/MS for global and phospho-proteomics

For phospho-proteomic studies control SH-SY5Y cells were transfected with Flag-WT-USP14 and cultured for 48h and lysed in IP-lysis buffer. Flag-WT-USP14 was immunoprecipitated utilizing Anti-Flag M2 (F1804, Sigma, Germany) antibody and eluted with 200 mM Glycine pH 2.0. Glycine- eluted samples were neutralized with 1.5 M Tris-HCl pH 8.4 and further processed for trypsin digestion and LC-MS analysis as below.

For global proteomic analysis, control and USP14-deleted cells were cultured for 48h, trypsinized and collected by centrifugation, followed by homogenization using lysis buffer containing 8 M urea, 100 mM ammonium bicarbonate, 2% sodium deoxycholate, 0.1% Octyl α-D -glucopyranoside, and five cycles of vortexing and bath sonication. Proteins were reduced and alkylated using tris (2-carboxyethyl) phosphine (TCEP) and iodoacetamide to a final concentration of 5mM and 50mM respectively and incubation in the dark for 30min.

10µg of protein were washed 8 to 10 times with 8M urea, 100mM ammonium bicarbonate in Amicon Ultra-0.5 centrifugal filters using a modified FASP method ^73^. Lysine-C endopeptidase solution (121-05063, FujiFilm Wako, Japan) in a ratio of 1:50 w/w was added to the protein lysates in about 4M urea/ 100mM ammonium bicarbonate and incubated overnight at room temperature (RT). The peptide digests were collected by centrifugation and trypsin solution was added in a ratio of 1:50 w/w in 50mM ammonium bicarbonate and incubated overnight at RT. The peptide samples were cleaned using Pierce™ C18 reverse-phase tips (Thermo Scientific, USA). Dried peptide digest was re-suspended in 0.3% TFA and sonicated in a water bath for 3min before injection. Digests were analyzed using nano-LC-MS/MS. The peptides were separated by Ultimate 3000 LC system (Dionex, Thermo Scientific, USA) equipped with a reverse-phase trapping column RP-2TM C18 trap column (0.075 x 10mm, Phenomenex, USA), followed by analytical separation on a bioZen C18 nano column (0.075 × 250 mm, 2.6 μm particle size; Phenomenex, USA). The injected sample analytes were trapped at a flow rate of 5µl x min-1 in 100% of solution A (0.1% formic acid). After trapping, the peptides were separated with a linear gradient of 125min comprising 110min from 3% to 35% of solution B (0.1% formic acid/80% acetonitrile), 6min to 45% B, and 4min to 95% of solution B. Each sample run was followed by an empty run to reduce the potential sample carryover from previous runs.

LC-MS acquisition was done using the mass spectrometer (Thermo Q Exactive HF) settings as follows: The resolution was set to 120,000 for MS scans, and 15000 for the MS/MS scans. Full MS was acquired from 350 to 1400 *m/z*, and the 15 most abundant precursor ions were selected for fragmentation with 45 s dynamic exclusion time. Ions with 2+, 3+, 4+, and 5+ charges were selected for MS/MS analysis. Maximum IT were set as 50 and 25ms and AGC targets were set to 3 e^6^ and 1 e^5^ counts for MS and MS/MS respectively. Secondary ions were isolated with a window of 1 *m/z* unit. Dynamic exclusion was set with a duration of 45sec. The NCE collision energy stepped was set to 28kJ mol–1.

### Proteomic data and bioinformatic analysis

Following LC-MS/MS acquisition, raw files were qualitatively analyzed by Proteome Discoverer (PD), version 2.5 (Thermo Scientific, USA). The identification of proteins by PD was performed against the human protein database (release 2023_1 with 20434 entries) using the built-in SEQUEST HT engine. The following parameters were used: 10ppm and 0.02 Da were mass error tolerance values set for MS and MS/MS, respectively. Trypsin was used as the digesting enzyme, and two missed cleavages were allowed. The carbamidomethylation of cysteine residues was set as a fixed modification, while the oxidation of methionine, deamidation of asparagine and glutamine, and phosphorylation of Serine and threonine were set as variable modifications. The false discovery rate was set to less than 0.01 and a peptide minimum length of six amino acids. Label-free quantification was done using unique peptides in Precursor Ion Quantifier. Differentially expressed proteins (DEPs) were identified based on the number of unique peptides used for label-free quantitation (≥2), at the FDR < 0.01 and the fold change (FC) from averaged, normalized protein intensities |≥1.5|, utilizing p≤ 0.05 by ANOVA in all comparisons, serving as inputs for Ingenuity Pathway Analysis (Qiagen™). The proteomics data were deposited to ProteomeXchange Consortium repository via MassIVE platform under the PXD047270 identifier.

### MitoTracker staining and Image analysis

Cells were cultured on µ-Slide 8-well ibiTreat plates (80806, Ibidi, Germany) and stained with 250nM MitoTracker DeepRed FM (Thermo Fisher Scientific, USA) for 30min at 37°C. Following the staining, the culture medium was replaced with FluoroBrite™ DMEM (Gibco, USA) imaging medium. The stained live-cells were imaged utilizing Nikon Eclipse Ti-E inverted widefield microscope with full environmental chamber, Hamamatsu Orca Flash 4.0 V2 B&W camera for fluorescence and Lumencor Spectra X light engine (Biomedicum Imaging Unit, Medicum, University of Helsinki).

Mitochondria branch length analysis was performed utilizing the ImageJ (NIH, Bethesda, USA) macro – Mitochondria Network Analysis (MiNA) ^74^. In each biological experiment, around 60-100 cells from different regions of the culture wells were analyzed for each sample and analyzed for differences in mean branch length parameter provided by MiNA. Briefly, pre-processing protocols provided by the macro were applied, and mitochondrial branches were skeletonized for analysis by MiNA. The changes in mitochondrial branch length measured as mean branch length parameter in the MiNA macro was utilized as a measure of change in mitochondria fusion/fission between control and USP14-deleted cells.

### CellROX Green staining

Cells were cultured and stained with 5µM CellROX green reagent (Thermo Fisher Scientific, USA) for 30min at 37 °C. Following the staining, the culture medium was replaced with FluoroBrite™ DMEM (Gibco, USA) imaging medium and imaged with EVOS FL cell imaging system (Thermo Fisher Scientific, USA).

For quantification of intensity, region of interest selection was made around each cell and intensity measured by the measure tool in ImageJ. Background intensity was measured and subtracted from the cell intensity to acquire a more accurate representation of CellROX staining intensity in the cell.

### Measurement of mitochondrial respiratory capacity

To study mitochondrial oxygen consumption rate (OCR), Control and USP14 deleted cells were plated on a Seahorse 96-well plate and cultured for 48h. As described before, basal Oxygen Consumption Rate (OCR) utilizing the Seahorse XFe96 analyzer (Seahorse Bioscience, Boston, MA, USA) ^75^ and the measurements were plotted as histogram.

### Transmission Electron Microscopy (TEM)

For thin-section TEM, Control and USP14 deleted cells were cultured for 48h on coverslips and fixed using 100 mM NaPO4, pH 7.4, 2% Glutaraldehyde (EM-grade, Sigma Aldrich, Germany) for 24hr, 4°C. Post-fixation, the cells were Osmicated utilizing Osmium tetroxide, dehydrated in ethanol series and acetone followed by gradual infiltration with Epon (TAAB 812) ^76, 77^. 60nm sections cut parallel to the coverslips were post-stained with uranyl acetate and lead citrate. Samples were then imaged utilizing the transmission electron microscope Jeol JEM-1400 electron microscope equipped with an Orius SC 1000B bottom-mounted charge-coupled device camera (Gatan, USA) at acceleration voltage of 80 kV.

### Statistical analysis

Depending on the experimental design, statistical analysis was performed utilizing Student’s t-test or one-way/two-way ANOVA. p-value, p < 0.05 was considered as statistically significant. Statistical analysis and the histogram plots were generated using GraphPad PRISM software.

## Conflict of interest

The authors declare no competing interests.

## Data availability

All the proteomics-related dataset has been submitted into MassIVE database under the PXD047270 identifier.

Uncropped blots for the IB images have been provided in supplementary.

## Acknowledgements

Supported by Liv och Hälsa Foundation, The Finnish Society of Sciences and Letters, Magnus Ehrnrooth Foundation, Finska Läkaresällskapet, Minerva Foundation. VS is supported by the Doctoral Programme in Biomedicine (DPBM), University of Helsinki. DL and Ml. are members of the COST-action: CA20113 - ProteoCure. LI and MK are supported by Centre of Excellence in Molecular Cell Engineering (Project No. 2014-2020.4.01.15-013), Estonia.

We thank Kristina Söderholm for skillful technical assistance. We are grateful to the following facilities for support: the proteomics analyses were performed at the Meilahti Clinical Proteomics Core Facility, HiLIFE supported by Biocenter Finland; EM-sample preparation and processing at EMBI, Institute of Biotechnology, University of Helsinki; the Nikon Eclipse Ti-E widefield inverted microscopy was done at Biomedicum imaging unit (BIU), HiLIFE, Medicum; gene mutations were done at Genome biology, Medicum and DNA sequencing was done at FuGU, University of Helsinki.

## Author contributions

DL and VS designed the project; VS, RS, LI, OE, NP, ML, MK, DL designed the experiments. VS, RS, LI, OE, NP, ML, MK, DL performed the experiments and analyzed the data. DL and VS wrote the manuscript; and all the authors have edited and approved the manuscript.

## Notes

### Competing Interest Statement

The authors have declared no competing interest.

