## Supplementary Figures 1-4 for "USP14 regulates pS129 α-synuclein levels and oxidative stress in human SH-SY5Y dopaminergic cells"

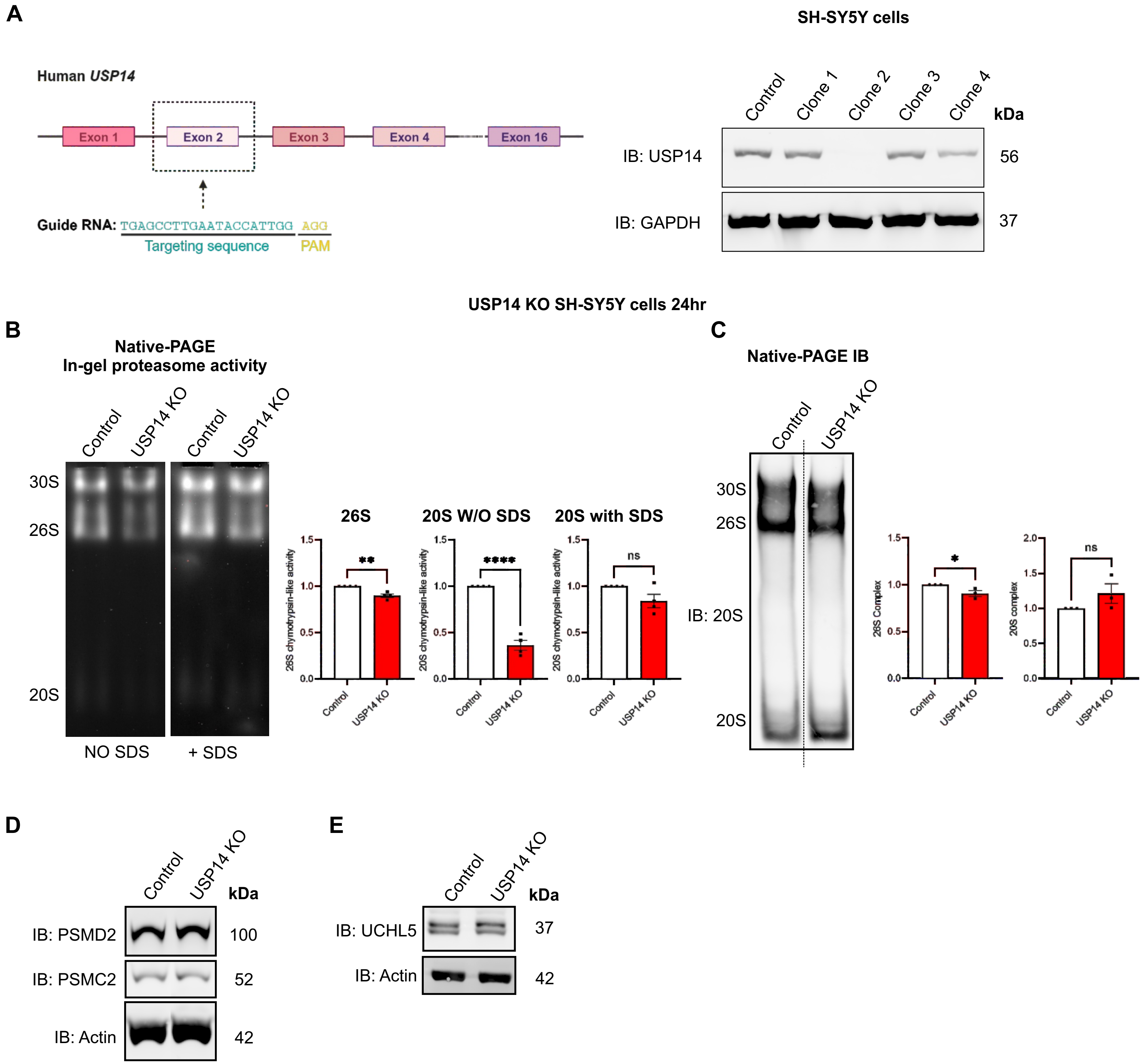
A
SH-SY5Y cells
Human USP14
Exon 1
Exon 2
Exon 3
Exon 4
Exon 16
Guide RNA: TGAGCCTTGAATACCATTGG AGG
Targeting sequence
PAM
Control
Clone 1
Clone 2
Clone 3
Clone 4
kDa
IB: USP14
56
IB: GAPDH
37
USP14 KO SH-SY5Y cells 24hr
B
Native-PAGE
In-gel proteasome activity
Control
USP14 KO
Control
USP14 KO
30S
26S
20S
NO SDS
+ SDS
26S
20S W/O SDS
20S with SDS
26S chymotrypsin-like activity
20S chymotrypsin-like activity
20S chymotrypsin-like activity
**
****
ns
1.5
1.0
0.5
0.0
Control
USP14 KO
C
Native-PAGE IB
Control
USP14 KO
30S
26S
IB: 20S
20S
26S Complex
20S complex
*
ns
2.0
1.5
1.0
0.5
0.0
Control
USP14 KO
D
Control
USP14 KO
kDa
IB: PSMD2
100
IB: PSMC2
52
IB: Actin
42
E
Control
USP14 KO
kDa
IB: UCHL5
37
IB: Actin
42

**CLEAR signaling pathway**

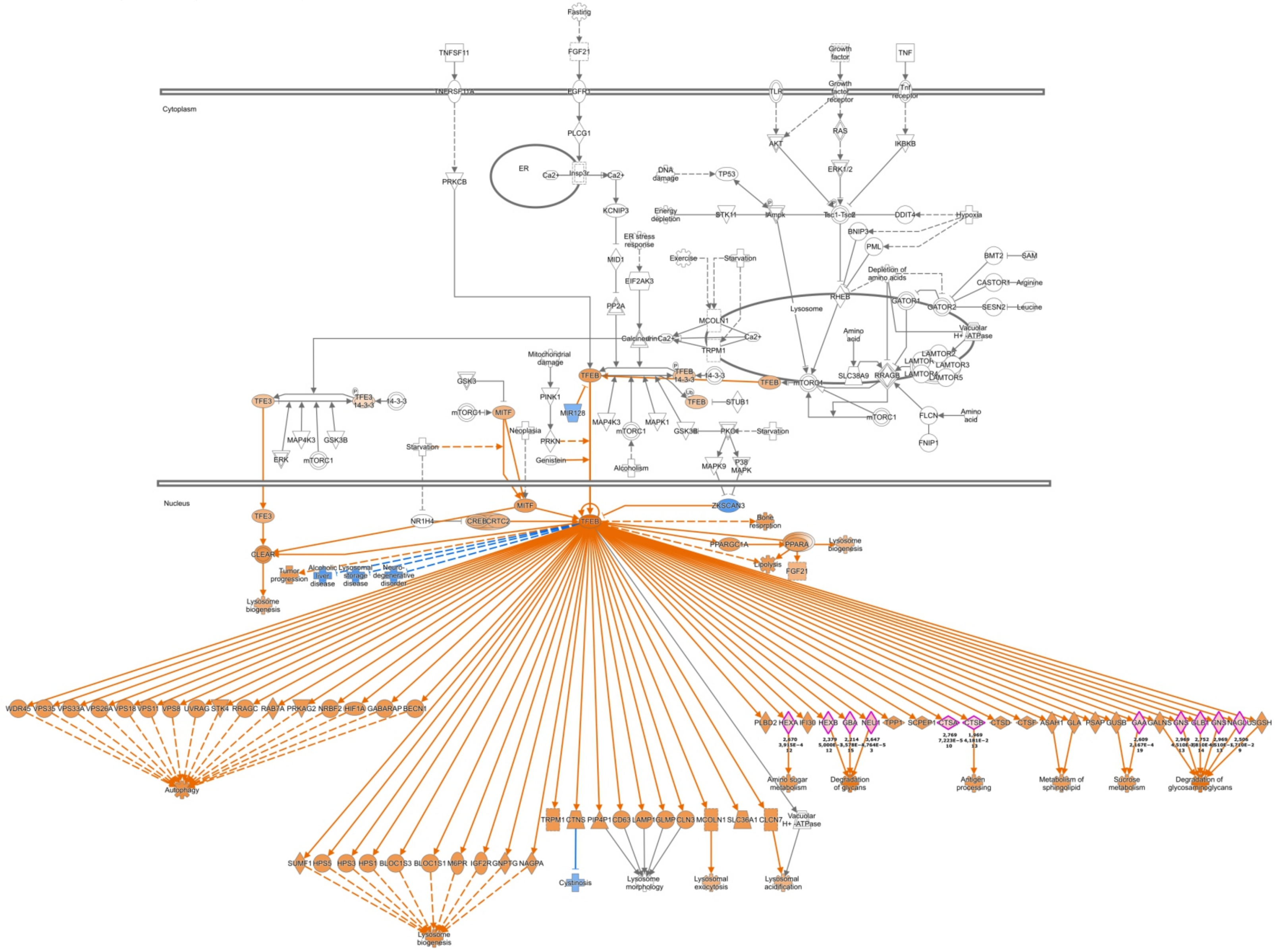


**Pathway:**

Increased activity

Decreased activity

No trend identified

Inconsistent finding

Direct effect

Indirect effect

Activator

Inhibitor

Identified DEP

**A** **USP14 KO cells**

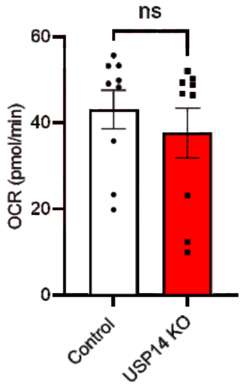


**B** **Figure 4C High exposure for OXPHOS cocktail IB**

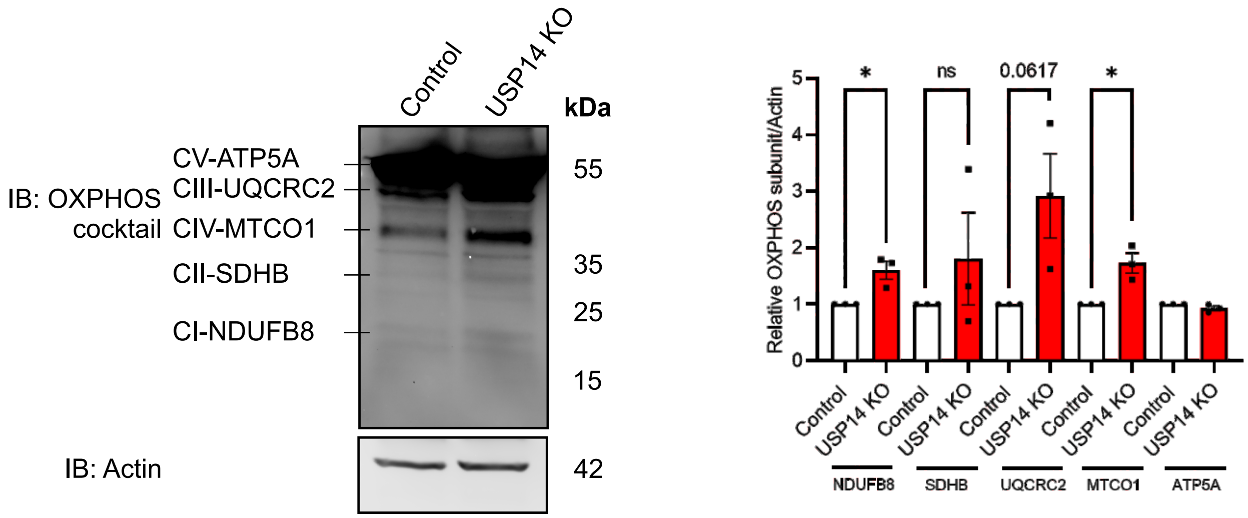


**C**

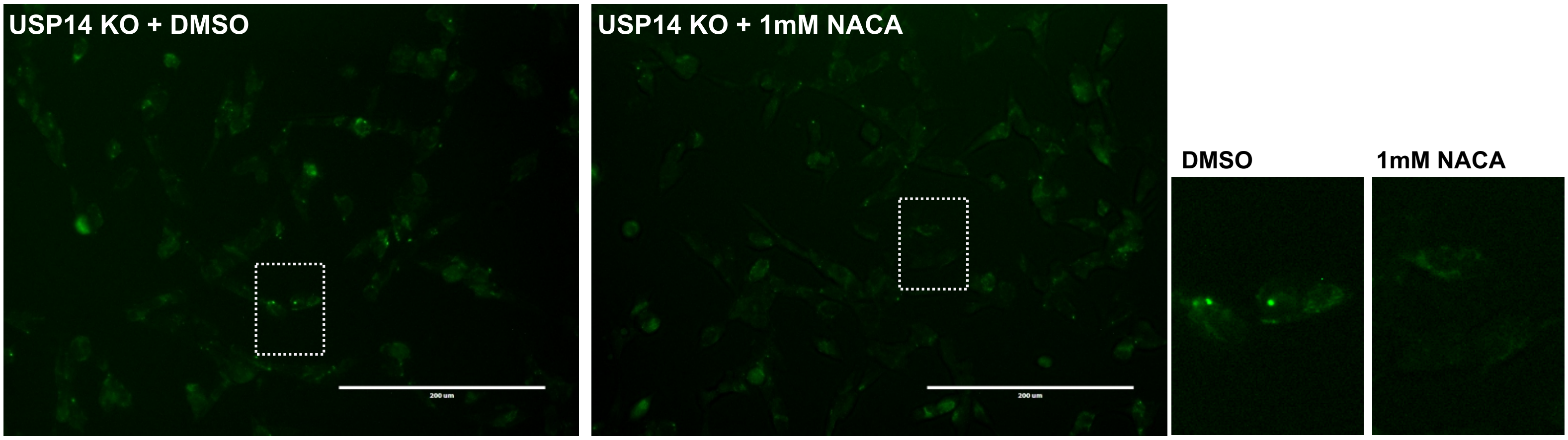

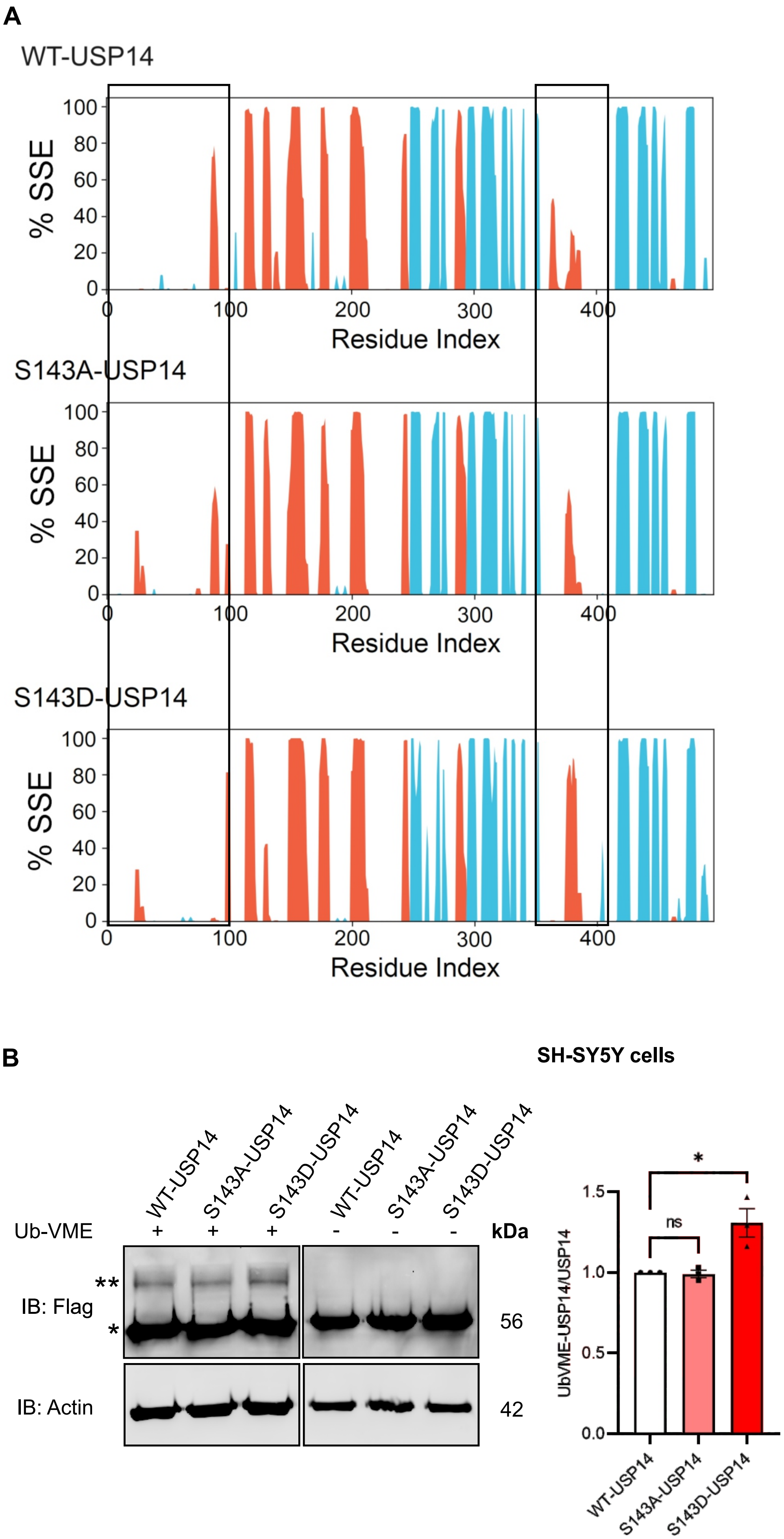
A
WT-USP14
% SSE
Residue Index
S143A-USP14
S143D-USP14
B
SH-SY5Y cells
WT-USP14
S143A-USP14
S143D-USP14
Ub-VME
kDa
IB: Flag
IB: Actin
56
42
UbVME-USP14/USP14
ns
*
